# Solaris: a panel of bright and sensitive hybrid voltage indicators for imaging membrane potential in cultured neurons

**DOI:** 10.1101/2024.02.02.578569

**Authors:** Junqi Yang, Siyan Zhu, Luming Yang, Luxin Peng, Yi Han, Rebecca Frank Hayward, Pojeong Park, Dachao Hu, Adam E. Cohen, Peng Zou

**Author notes:** Correspondence (P.Z.). These authors contributed equally (Junqi Yang, Siyan Zhu).

## Abstract

Dynamic changes in the membrane potential underlie neuronal activities. Fluorescent voltage indicators allow optical recording of electrical signaling across a neuronal population with cellular precision and at millisecond-level temporal resolution. Here we report the design and characterization of a chemigenetic hybrid voltage indicator, Solaris, in which a circularly permuted HaloTag is inserted into the first extracellular loop of *Acetabularia* rhodopsin. Solaris is compatible with fluorogenic HaloTag ligands JF_525_, JF_549_, JF_552_, JF_585_, and JF_635_. The most sensitive conjugate, Solaris_585_, has more than 2-fold higher voltage sensitivity than the spectrally similar Voltron2_585_ (ΔF/F_0_ = -28.1 ± 1.3% versus -12.3 ± 0.7% per action potential in cultured neurons). Solaris_585_ supports the measurement of optogenetically evoked spike activity or dual-color imaging in conjunction with green-emitting calcium or glutamate indicators. Solaris indicators are also applicable to fluorescence lifetime imaging, which probes the absolute membrane potential. This new hybrid voltage indicator is a valuable tool for imaging neuronal electrophysiological activities in cultured cells with substantially improved dynamic range compared to previous hybrid indicators.

## Introduction

Fluorescent voltage indicators have pushed the non-invasive detection of electrical signaling in neurons with unprecedented spatiotemporal resolution^1-3^. To achieve high signal-to-noise ratio (SNR) detection of neuronal action potentials (APs), an ideal voltage indicator should have: (1) high voltage sensitivity across the physiological range of -80 mV to 40 mV; (2) sub-millisecond responses to voltage changes; (3) bright fluorescence to yield sufficient photons at kilohertz imaging rate; and (4) low photo-toxicity and high photostability to allow long-time voltage imaging (i.e. >30 min). Purely protein-based genetically encoded voltage indicators (GEVIs) are often hampered by low brightness and/or poor photostability, which are intrinsic properties of fluorescent proteins (FPs) as compared to synthetic fluorophores^4-8^.

The recent development of chemigenetic hybrid voltage indicators has combined the superior photophysical properties of organic dyes and the genetic targetability of GEVIs. Although microbial rhodopsin-based GEVIs have a highly sensitive response to membrane voltage changes within a millisecond (ΔF/F_0_ = 48 ± 3% for QuasAr2 for a single AP)^9^, they are often hampered by poor fluorescence quantum yields (e.g. Archon1 has QY ≈ 0.49%)^10^. Electrochromic fluorescence resonance energy transfer (eFRET) provides a way to improve the brightness of rhodopsin-based GEVIs while maintaining their fast kinetics. In eFRET indicators, a bright organic fluorophore^4-7,11^ or a fluorescent protein^12-14^ acts as the fluorescent reporter. As the absorption spectrum of rhodopsin (acceptor) is modulated by cellular membrane potential, the emission of the appended fluorophore (donor) changes. For example, in Voltron indicators, the voltage-sensing *Acetabularia acetabulum* rhodopsin mutant (Ace) is fused with the self-labeling HaloTag protein that covalently binds the ligand modified with a synthetic fluorophore. When labeled with bright and stable rhodamine dyes such as JaneliaFluor (JF) dyes, the resulting chemigenetic hybrid indicators allow sensitive detection of AP spikes in mouse, zebrafish, and fruit fly^4,5^. The more recently developed Voltron2_525_ responds to single APs with ΔF/F_0_ = -10.1 ± 1.5%^5^.

Our group recently reported the development of hybrid voltage indicators (HVIs) that exhibit high voltage sensitivity (ΔF/F_0_ = -25.3 ± 0.8% per AP for HVI-Cy3). HVIs contain a small-molecule dye that is site-specifically linked to a voltage-sensing rhodopsin protein via enzyme-mediated probe-ligation and bioorthogonal conjugation^6^. Due to the smaller tag size compared to the HaloTag receptor, the fluorescent dye reporter is positioned closer to the retinal chromophore, leading to higher FRET efficiency and thus higher voltage sensitivity than in the Voltron family. Indeed, the dynamic range of HVI-Cy3 is more than two-fold higher than the spectrally similar Voltron2_525_. To further improve the photostability and to reduce phototoxicity, cyclooctatetraene (COT)-modified cyanine dyes have been utilized in HVIs^7^. HVI-COT dyes have enabled high-fidelity continuous voltage imaging in cultured neurons for longer than 30 min. However, HVIs are not directly applicable to *in vivo* imaging due to the requirement of enzymatic labeling.

Here, we report the design and characterization of a novel hybrid voltage indicator, with a circularly permuted HaloTag receptor (cpHaloTag) inserted into the first extracellular loop (ECL1) of Ace rhodopsin protein. The resulting indicator, termed Solaris, can be conjugated with diverse fluorogenic HaloTag ligands, including JF_525_, JF_549_, JF_552_, JF_585_, and JF_635_. Solaris faithfully reports APs with more than 2-fold higher voltage response than Voltron2. The compatibility of Solaris with red or far-red dyes allows multiplexed imaging with blue-shifted optogenetic actuators, such as CheRiff^9^ or green fluorescent calcium indicators, such as GCaMP6s^15^. Current versions of Solaris traffic well in cultured cells but not *in vivo*. Efforts are underway to improve the trafficking *in vivo*.

### Design and characterization of novel hybrid voltage indicator

We recently developed a pair of red eFRET GEVIs called Cepheid1b (featuring high brightness) and Cepheid1s (featuring high photostability)^16^ that fused red fluorescent proteins at the ECL1 of Ace. Cepheid1b/s exhibited improved voltage response, brightness, and photo-stability, and have been successfully applied to imaging voltage dynamics in mouse brain slices. We sought to combine the high voltage sensitivity of Cepheid with the brightness and spectral tunability of HaloTag-based hybrid voltage indicators. We replaced the fluorescent protein in ECL1 with a circularly permuted HaloTag (cpHaloTag) (Fig. 1A). To improve membrane trafficking, we also fused triple-tandem transport signal (3xTS), ER-exiting (ER2) sequences^17^ and dark mOrange2 at the C-terminus of the rhodopsin^6,7,16^. We termed the resulting construct Solaris.

**Figure 1.**
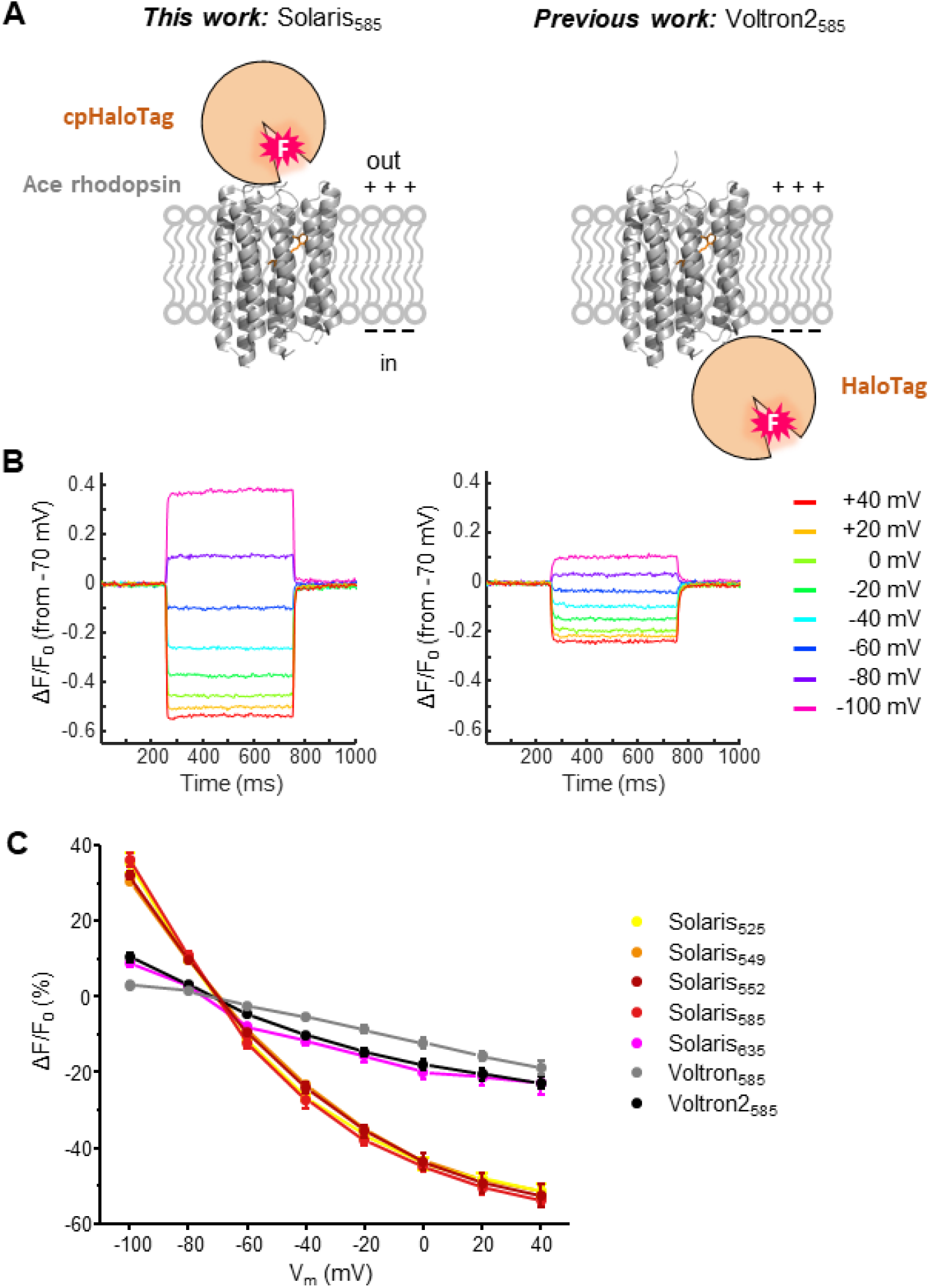
Design and characterization of Solaris voltage indicator. **(A)** Scheme of predicted structures of Solaris and Voltron2. **(B)** Comparing voltage sensitivities of Solaris_585_ (left) and Voltron_2585_ (right) in HEK293T cells. **(C)** Normalized fluorescence-voltage response curves of Solaris_525_, Solaris_549_, Solaris_552_, Solaris_585_, Solaris_635_, Voltron_585_ and Voltron_2585_ in HEK293T cells.

We labeled Solaris in human embryonic kidney 293T (HEK293T) cells with a variety of dyes, with emission peaks ranging from approximately 550 nm to 650 nm (JF_525_, JF_549_, JF_552_, JF_585_ and JF_635_). To measure voltage sensitivity, we applied whole-cell voltage clamp to vary the membrane potential stepwise in increments of 20 mV. Among the above dyes, Solaris_585_ exhibited the highest voltage response when the membrane potential was changed from -80 mV to 20 mV, with whole-cell fluorescence changing by ΔF/F_0_ = -60.6 ± 1.6% (mean ± s.e.m., n = 5 cells, illuminated with 594 nm laser at 1.72 W/cm^2^) (Fig. 1B). Solaris labeled with other dyes had voltage sensitivities ranging between ΔF/F_0_ = -58% to-24% per 100 mV (Fig. 1C). The voltage sensitivity of Solaris compares favorably to Voltron series. For example, the fluorescence of Voltron_585_ and Voltron2_585_ changed only by -17.1 ± 1.3% (n = 6 cells) and -24.0 ± 1.5% (n = 7 cells), respectively, for the same range of depolarization (Fig. 1C). At room temperature, the response half-time of Solaris_585_ was *τ*_1/2_ = 1.75 ± 0.11 ms in the depolarizing step (n = 5 cells, acquired at camera frame rate of 1058 Hz), which was slightly faster than Voltron2_585_ (*τ*_1/2_ = 2.1 ± 0.1 ms, n = 5 cells), and comparable to Cepheid1b/s (*τ*_1/2_ = 1.9 ± 0.1 ms and 1.5 ± 0.2 ms, respectively).

We also compared the voltage sensitivities of Solaris and Voltron2 in cultured rat hippocampal neurons. Solaris labeled with different dyes exhibited good expression and membrane localization in the soma and neurites of cultured neurons (Fig. 2A). To induce the firing of individual APs, we injected a current of 200-400 pA into cultured neurons for a duration of 5-10 ms, with a repetition rate of 2-4 Hz. Consistent with the measurement of F-V curve in HEK293T cells, Solaris_585_ has the highest voltage sensitivity when reporting APs, with ΔF/F_0_ more than 2-fold higher than Voltron2_585_ (-28.1 ± 1.3% vs. -12.3 ± 0.7%, n = 7 cells each) when measured under matched conditions (Fig. 2B).

**Figure 2.**
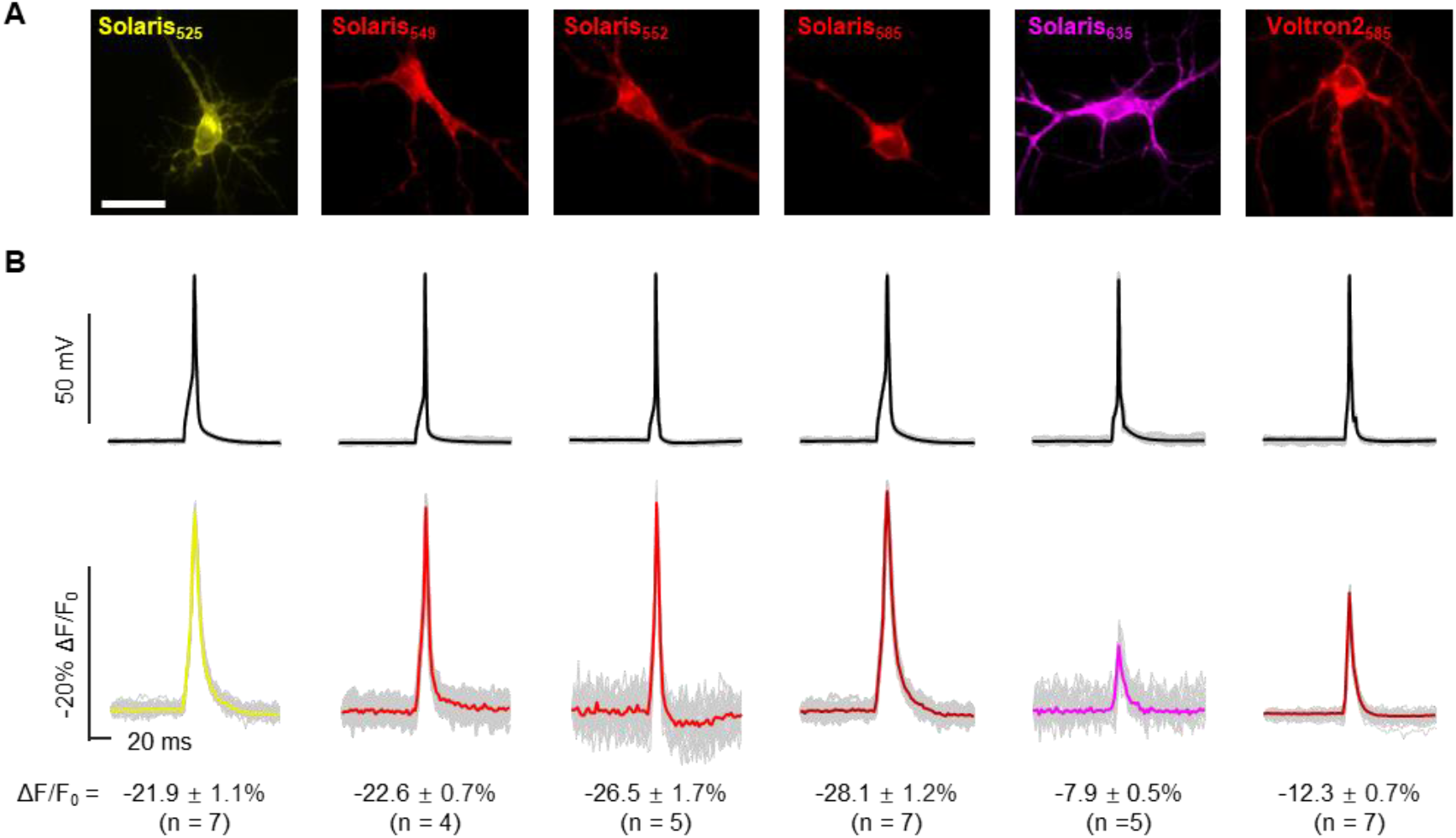
Voltage imaging with Solaris in cultured rat hippocampal neurons. **(A)** Images of rat hippocampal neurons expressing Solaris labeled with different dyes, and Voltron_2585_. Scale bar, 20 μm. **(B)** Electrophysiological and corresponding optical waveforms recorded from neurons expressing Solaris labeled with different dyes or Voltron_2585_. Stimulated AP firing was triggered via current injection into rat hippocampal neurons. Twenty successive stimuli (grey) are superimposed, and the averaged traces of fluorescence are shown in colors.

### Multiplexed voltage imaging with Solaris

The red-shifted spectra of Solaris_585_ allow the combination with the blue-light-activated cation channel CheRiff for all-optical electrophysiology. In cultured rat hippocampal neurons co-expressing CheRiff and Solaris_585_, we applied a 488 nm laser to stimulate AP firing while imaging membrane voltage simultaneously (Fig. 3A). Solaris_585_ could faithfully record APs under various blue-light-activate patterns (Fig. 3B).

**Figure 3.**
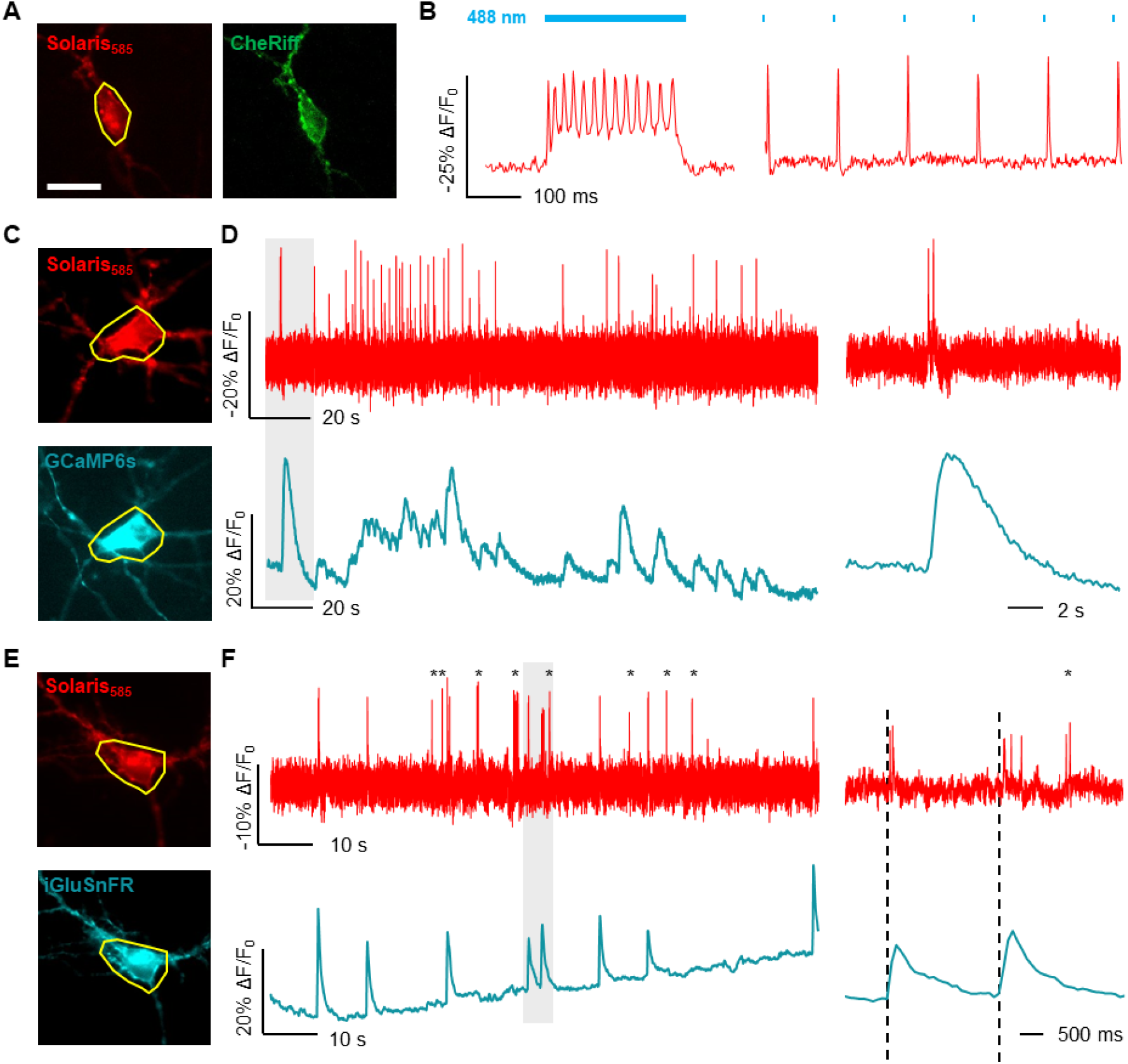
All-optical electrophysiology and multiplexed imaging with Solaris_585_. **(A)** Images of rat hippocampal neurons expressing CheRiff-EGFP and Solaris_585_. **(B)** Fluorescence of Solaris_585_ in response to optogenetically triggered AP firing. Neurons expressing Solaris were imaged under 561 nm wide-field illumination at 6.60 W/cm^2^ and simultaneously stimulated with a pulsed 488 nm laser at 0.41 W/cm^2^. Image series were acquired at a camera frame rate of 400 Hz. **(C)** Images of cultured rat hippocampal neuron co-expressing GCaMP6s-NES and Solaris_585_. **(D)** Dual-color imaging of calcium (green) and voltage (red) activities in cultured neurons, with a zoom-in view of the shaded regions shown on the right. **(E)** Images of a cultured rat hippocampal neuron co-expressing SF-iGluSnFR and Solaris_585_. **(F)** Dual-color imaging of extracellular glutamate (green) and voltage (red) activities in cultured neurons, with a zoom-in view of the shaded region shown on the right. Co-expression for all three pairings was achieved via Lipofectamine 3000 transfection.

Furthermore, we applied Solaris_585_ with green-emitting indicators to monitor membrane potential simultaneously with other neuronal signals such as cytoplasmic calcium or extracellular glutamate. In neurons co-expressing Solaris_585_ and the green calcium indicator GCaMP6s-NES (Fig. 3C), matched calcium spikes and AP spikes could be observed (Fig. 3D).

We also paired Solaris_585_ with SF-iGluSnFR to record membrane potential and glutamate release (Fig. 3E). The occurrence of the extracellular glutamate events often coincided with the firing of AP spikes (Fig. 3F). However, as marked by asterisks in Fig. 3F, there were also instances where AP spikes did not coincide with glutamate signals around the soma, presumably reflecting excitatory inputs outside the field of view. Taken together, the above results demonstrate that Solaris_585_ has the ability for multiplexed imaging of membrane potential with other physiological signals.

### Fluorescence lifetime imaging with Solaris

While intensiometric voltage imaging could report relative changes in the membrane potential (e.g. action potential spikes and sub-threshold depolarizations), it is often challenging to determine the absolute value of membrane potential (e.g. values of resting membrane potential). Indeed, intensiometric recording is susceptible to variations in fluorophore concentration, photobleaching, illumination intensity and collection efficiency, background autofluorescence and cellular morphology. Fluorescence lifetime imaging (FLIM) provides an alternative method for accurate quantification of membrane potential, because the fluorescence lifetime is an intrinsic property of the fluorophore that is insensitive towards the above factors.

FLIM measurement has been previously applied to voltage imaging to determine absolute membrane potential. Brinks *et al*. demonstrated that eFRET indicator CAESR exhibits voltage-dependent changes in the lifetimes under two-photon illumination^18^. Lazzari-Dean *et al*. used FLIM with the photo-induced electron transfer (PeT) dye VF2.1Cl to track membrane potential changes with an accuracy of 5 mV and reported membrane potential distributions over thousands of cells^19^. However, indicators used in these studies have relatively small changes in lifetime over physiological voltage ranges. For example, CAESR and VF2.1Cl respond to 100 mV membrane potential change with Δτ/τ_0_ = -6.1 ± 0.8% and 22.4 ± 0.4%, respectively^19^.

The eFRET voltage sensing mechanism of Solaris indicates that voltage fluctuations will modulate the energy transfer efficiency of the appended fluorophore, thus changing its fluorescence lifetime. To measure the voltage sensitivity of fluorescence lifetime, we constructed a HEK293T cell line stably expressing an inward rectified potassium channel Kir2.1. Due to the action of Kir2.1, the resting membrane potential of this cell line is lowered to -73.7 ± 3.1 mV (n = 7 cells, mean ± s.e.m.). When Kir2.1-expressing cells were treated with the channel-forming ionophore Gramicidin A at 2 μg/mL for 10 min, the membrane potential was depolarized to 2.2 ± 1.0 mV (n = 6 cells), as measured by whole-cell patch-clamp recording (Fig. 4A).

**Figure 4.**
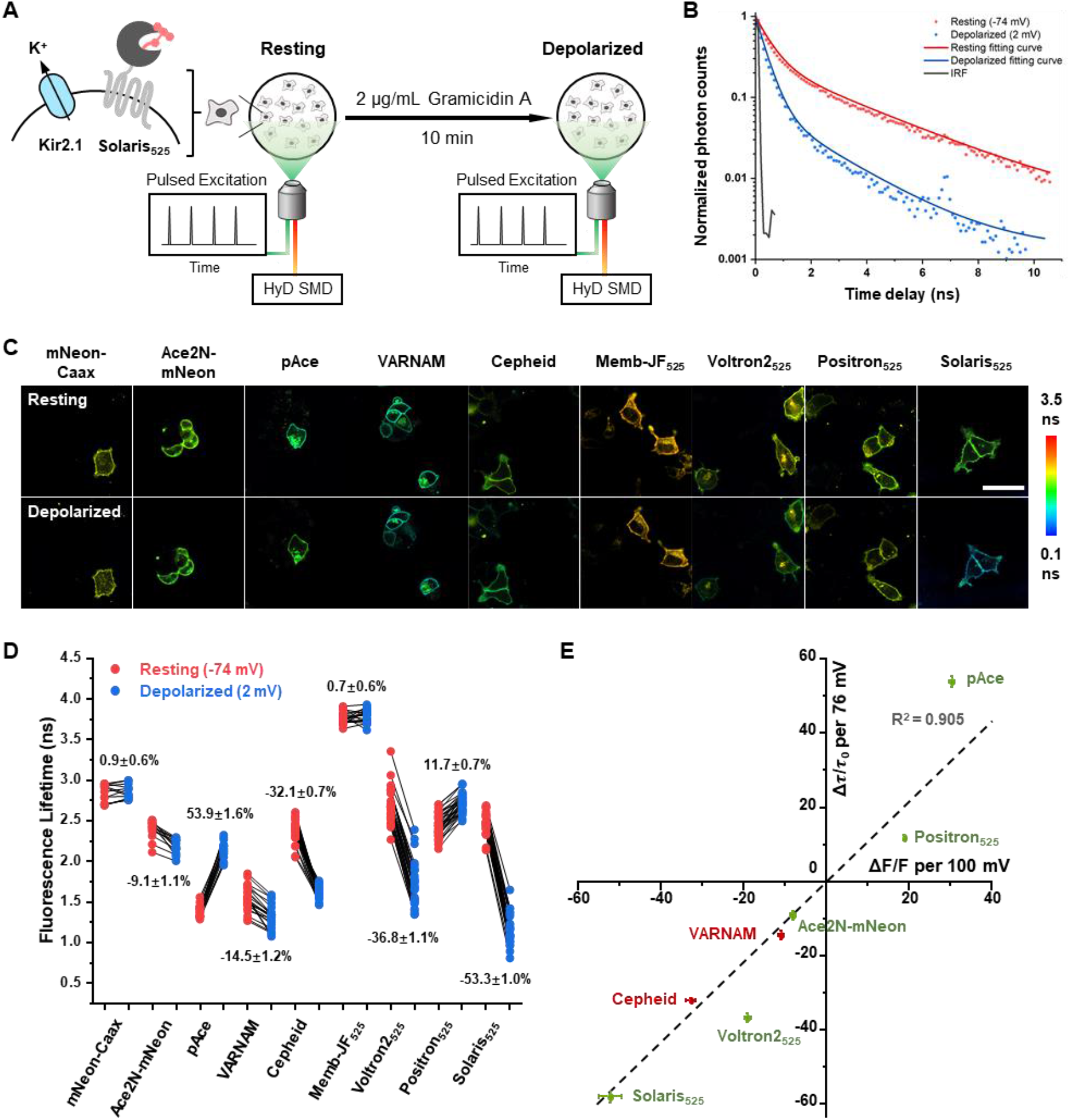
Fluorescence lifetime imaging with Solaris. **(A)** Schematic of fluorescence lifetime measurement with time-correlated single-photon counting (TCSPC) in HEK293T cells expressing Kir2.1. **(B)** Fluorescence lifetime decay and fitting curve of Solaris_525_ in resting and depolarized states. **(C)** Representative fluorescent lifetime images of a series of GEVIs in resting and depolarized states. Voltron2, Positron and Solaris were labeled with JF_525_. Scale bar, 50 μm. **(D)** Statistical analysis of voltage sensitivity in fluorescence lifetime for a panel of GEVIs. Each dot represents measurement on a single cell, numbers indicate mean ± s.e.m. **(E)** Correlation between voltage sensitivities measured in fluorescence lifetime (Δτ/τ_0_) versus fluorescence intensity (ΔF/F).

We applied the above depolarization method to calibrate the voltage sensitivity of a series of eFRET GEVIs under single-photon time-correlated single-photon counting (TCSPC) FLIM, including Ace2N-mNeon^12^, pAce^14^, VARNAM^13^, Cepheid^16^, Voltron_525_ ^5^, Positron_525_ ^20^ and Solaris_525_ (Fig. 4B-C). In negative control samples expressing voltage-insensitive reporters, such as membrane-anchored mNeonGreen (mNeon-Caax) and membrane-anchored HaloTag-JF_525_ (Memb-JF_525_), little change in fluorescence lifetime (<1.0%) was observed upon membrane depolarization. In contrast, all GEVIs exhibited changes in lifetime upon membrane depolarization, with Solaris_525_ showing the highest sensitivity of Δτ/τ_0_ = -53.3 ± 1.0% (n = 29 cells, mean ± s.e.m.), which was 1.5-fold and 4.5-fold higher than Voltron2 (-36.8 ± 1.1%, n = 29 cells) and Positron (11.7 ± 0.7%, n = 35 cells), respectively (Fig. 4D). Notably, the voltage sensitivities measured in lifetime recording (Δτ/τ_0_) were highly correlated with the voltage sensitivities measured in intensiometric recording (ΔF/F), as expected from the eFRET voltage sensing mechanism (Fig. 4E).

## Conclusions

In summary, we have reported a panel of fast and sensitive genetically targetable voltage indicators named Solaris. Among them, Solaris_585_ features a high voltage sensitivity of -53% ΔF/F_0_ per 100 mV, which is more than twice as sensitive as Voltron2_585_ (-22% ΔF/F_0_ per 100 mV). The high dynamic range of Solaris_585_ allows sensitive detection of APs at 484 Hz frame rate in cultured neurons (ΔF/F_0_ = -28%) under illumination of 594 nm laser. Solaris_585_ supports multiplexed imaging with green fluorescent indicators and all-optical electrophysiology. Solaris_525_ also shows high sensitivity with FLIM. We envision that the sensitivity of Solaris indicators could be further optimized through directed evolution.

Current versions of Solaris traffick well in cultured neurons but not *in vivo*. We speculate that the insertion of a protein tag into the extracellular loop of Ace has destabilized the protein structure. This may be improved through linker optimization, an effort that is underway in our laboratory.

## Methods

### Molecular cloning

Plasmids were generated by Gibson Assembly. PCR amplified the inserts and the vector with a 15-bp overlap. These DNA fragments were mixed with Gibson Assembly enzyme (Lightning). The sequences of plasmids were verified by sequencing. The plasmids and DNA sequences are available from the corresponding author upon request.

|  | Vector | Promoter | Sequence features |
| --- | --- | --- | --- |
| Solaris | FCK | CaMKII | Ace(1-68)-cpHaloTag-Ace(69-227)(S81)-TS-mOrange2(Dark)-ER2 |
| Voltron | FCK | CAMKII | Ace2N-HaloTag-TS-ER2 |
| Voltron2 | FCK | CAMKII | Ace2N(D122)-HaloTag-TS-ER2 |

### Expression of Solaris in cultured cells

HEK293T cells were incubated in Dulbecco’s Modified Eagle medium (DMEM, Gibco) containing 10% v/v fetal bovine serum (FBS, Gibco) at 37°C with 5% CO_2_. For transfection, cells were plated in a 24-well plate and grown to 70-90% confluent. For each well, 500 ng plasmid and 1 μL Lipofectamine 2000 (Invitrogen) reagent were mixed in Opti-MEM medium and incubated for 15 min at room temperature. The mixture was added to DMEM and incubated with the cells for 4-6 h. Thereafter, cells were treated with trypsin-EDTA (0.05%, Gibco) for 1 min at room temperature, replated onto a 14-mm matrigel-coated glass coverslip at 10% dilution, and then incubated in complete medium for 24 h before imaging.

Primary rat hippocampal neurons were digested from dissected rat brains at postnatal day 0 (P0) and seeded on 14-mm glass coverslips pre-coated with 20 μg/mL poly-D-lysine (Sigma) and 10 μg/mL laminin (Gibco). Neurons were incubated at 37 °C with 5% CO_2_ and were transfected on DIV8 (8 days *in vitro*). 250-500 ng plasmid and 1 μL Lipofectamine 3000 reagent (Invitrogen) were mixed in Neurobasal medium for a well and incubated for 15 min at room temperature. Neurons were then incubated with the mixture for 45 min at 37 °C with 5% CO_2_. Transfected neurons were imaged after 3-7 days.

### Fluorescence voltage imaging in HEK293T cells and cultured neurons

All of the fluorescence imaging experiments were conducted on an inverted microscope (Nikon-TiE) equipped with a 40x, NA1.3 oil immersion objective lens, five laser lines (Coherent OBIS 488 nm, 532 nm, 561 nm, 594 nm and 637 nm), and a sCMOS camera (Hamamatsu ORCA-Flash4.0 v2). The microscope, lasers and cameras were controlled by a custom-built software written in LabVIEW (National Instruments, 15.0 version).

For JaneliaFluor (JF) dye labeling, transfected HEK293T cells or neurons were incubated in Dulbecco’s modified Eagle medium (DMEM, Gibco) with 100 nM JF dyes for 15 min at 37 °C. After that, cells were gently rinsed three times with DMEM. The labeled HEK293T cells or neurons on coverslips were transferred to Tyrode’s buffer before imaging. Image analysis was performed in MATLAB (version R2018b) and ImageJ/Fiji (version 1.53t).

A dual-view device (Photometrics DV2, with a dichroic mirror T565lpxr and two emission filters of ET525/50m and ET610LP (Chroma)) was used to split the emission fluorescence into green/red fluorescence channels for two-color simultaneous imaging. For simultaneous imaging of membrane voltage and calcium, GCaMP6s-NES and Solaris were co-transfected into neurons at DIV8. On DIV11-15, neurons were illuminated with 488 nm (0.41 W/cm^2^) and 561 nm (6.60 W/cm^2^) lasers and imaged with a dual-view device at a frame rate of 400 Hz. For simultaneous imaging of membrane voltage and glutamate, iGluSnFR and Solaris were co-transfected into neurons on DIV8 and imaged on DIV11-15 in dual-view mode with 488 nm (0.41 W/cm^2^) and 561 nm (6.60 W/cm^2^) lasers at a frame rate of 400 Hz. The traces of GCaMP6-NES and SF-iGluSnFR in Figure 3 were displayed after downsampling to 10 Hz.

### Electrophysiology for cultured cells

To measure the dynamic range and kinetics of the voltage indicators in HEK293T cells, the membrane potential was recorded and clamped with whole-cell voltage clamp (Axopatch 200B, Axon Instruments). Borosilicate glass electrodes (Sutter) were pulled to be patch pipettes with tip resistances of 4-8 MΩ. The intracellular (IC) solution contained 125 mM potassium gluconate, 8 mM NaCl, 1 mM EGTA, 10 mM HEPES, 0.6 mM MgCl_2_, 0.1 mM CaCl_2_, 4 mM Mg-ATP and 0.4 mM GTP-Na_2_ (295 mOsm/kg, pH 7.3). The extracellular (XC) solution for patch clamp recording contained 125 mM NaCl, 2.5 mM KCl, 1 mM MgCl_2_, 3 mM CaCl_2_, 10 mM HEPES and 30 mM glucose (305-310 mOsm/kg, pH 7.3) at room temperature. For patch clamp measurement of HEK293T cells, the XC solution was supplemented with 50 μM 2-APB to block gap junction. Fluorescence image series were recorded at the frame rate of 1058 Hz.

To measure the response of Solaris to action potentials, cultured rat hippocampal neurons at DIV11-15 were current-clamped and injected with 200-300 pA of current for 5-10 ms to stimulate action potential firing. Fluorescence imaging of neurons was performed at the camera frame rate of 484 Hz.

### All-optical electrophysiology

All-optical electrophysiology experiments were conducted on cultured hippocampal neurons co-expressing CheRiff and Solaris. Co-expression was achieved via Lipofectamine 3000 transfection. Transfected neurons were stimulated with repeated 2 ms (0.08 W/cm^2^) 488 nm light pulses, or repeated 250 ms (0.04 W/cm^2^) 488 nm light step. Images were captured at a frame rate of 400 Hz under constant 594 nm illumination (1.72 W/cm^2^).

### Fluorescence lifetime imaging experiments

FLIM imaging was performed on a Leica TCS SP8 STED 3X inverted scanning confocal microscope (Leica Microsystems, Germany) equipped with a fluorescence lifetime imaging module (Leica SP8 FALCON). White-Light laser was supplied by NKT Photonics (SuperK EXTREME EXW-12, NIM) and provided 80 MHz pulsed excitation with wavelength of 470 to 670 nm. Fluorescence emission was collected through HCX PL APO 100x/1.4 STED oil immersion objective (Leica). Emitted photons were detected with a hybrid detector HyD SMD (Leica). The detector was cooled by a recirculating water chiller (Laird, THERMAL SYSTEMS). Data were acquired with a scan speed of 400 Hz and 1024 × 1024 pixels. The laser excitation wavelength and fluorescence emission bandwidth are summarized below:

|  | Ex./nm | Em./nm |
| --- | --- | --- |
| mNeon-Caax | 488 | 510-600 |
| Ace2N-mNeon | 488 | 510-600 |
| pAce | 488 | 510-600 |
| VARNAM | 561 | 510-600 |
| Cepheid | 637 | 650-700 |
| memb-JF<sub>525</sub> | 525 | 550-650 |
| Voltron-JF<sub>525</sub> | 525 | 550-650 |
| Positron-JF<sub>525</sub> | 525 | 550-650 |
| Solaris-JF<sub>525</sub> | 525 | 550-650 |

Fluorescence lifetime decay data were fitted using LAS X (Leica), which solved the non-linear least square problem using the Levenberg-Marquadt algorithm. By adjusting the number of exponential decay components to achieve the best fitting effect, Solaris_525_ lifetime data were fit to a sum of two exponential decay components (Equation 1). Attempts to fit the Solaris_525_ data with a single or two exponential decay were unsatisfactory.

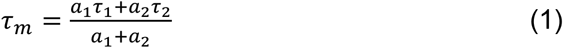

The fluorescence lifetime of Memb-JF_525_ was adequately described by a single exponential decay (Equation 2).

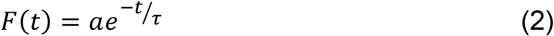

### Data analysis and statistics

Both the electrical and optical data were analyzed with custom software written in MATLAB (MathWorks, version R2018b). Fluorescence traces shown in Figs. 1 and 2 were derived from averaged movie sequences. The fluorescence traces in Fig. 1B were generated by averaging the imaging data series corresponding to 30 periods of square-wave voltage-clamp command signals over about 12 s. The fluorescence traces in Fig. 2B were derived as the spike-triggered average movies by aligning and averaging 40 AP spikes evoked by current injections over about 10 s. For waveform comparison and sensitivity calculation (Figs. 1 and 2), traces were extracted from regions-of-interest (ROIs) manually selected on the membrane of HEK293T cells (Fig. 1B) or the soma of neurons (Figs. 1D and 2E).

For multiplex imaging of voltage and Ca^2+^/glutamate signals, fluorescence traces of two channels were extracted directly from the ROIs selected around the soma of the neuron (Fig. 3, outlined in yellow). Prior to averaging, photobleaching removal was applied to every voltage imaging trace after background subtraction, and the Ca^2+^/glutamate traces were downsampled to 10 Hz from the original frequency of 400 Hz.

For the resting potential measurement of HEK293T cells expressing Kir2.1 channel with or without treatment of 2 μg/mL Gramicidin A (Fig. 4B), cells were held at -80 mV, and the resting potential was determined by the membrane potential value at I = 0 mode when the membrane was just ruptured and the whole-cell current clamp was established.

